# Nanopore sequencing of nested nrDNA barcodes reliably identifies orchid bees (Euglossini, Apidae)

**DOI:** 10.64898/2026.08.25.747120

**Authors:** Andreas Kolter, Mabel Alvarado, David W Roubik, Thomas Eltz

**Affiliations:** Department of Biology, Genetics, Ecology and Evolution, Aarhus University, Denmark; Centre for Biodiversity Genomics, Department of Integrative Biology, College of Biological Science, University of Guelph, Guelph, Canada; Departamento de Entomología, Museo de Historia Natural, Universidad Nacional Mayor de San Marcos, Lima, Peru; Smithsonian Tropical Research Institute, Panama, emeritus; Department of Animal Ecology, Evolution and Biodiversity, Faculty of Biology and Biotechnology, Ruhr University Bochum, Bochum, Germany

## Abstract

Orchid bees (Euglossini, Apidae) are Neotropical insects whose species-level identification can depend on minute morphological characters, some difficult to see or analyze. In such cases, DNA barcoding may facilitate identification by comparing standardized DNA sequences with reference libraries. The mitochondrial cytochrome c oxidase I (COI) marker widely used in animals does not, however, provide uniform species-level resolution across bee lineages. We developed an adaptive-length nuclear ribosomal DNA (nrDNA) barcoding framework based on overlapping Nanopore-sequenced markers spanning approximately 500 to 5500 bp for 114 Euglossini species. By matching barcode length to specimen quality, material with varied preservation histories was processed within a single analysis. Leave-k-out validation with IDTAXA achieved more than 96% identification success for two longer barcodes, while performance was lower for the shortest. Combining barcode lengths within one reference library maintained high identification success, and confidence filtering reduced overclassification when species were absent from the reference library. For orchid bees, this framework permits affordable high-throughput identification and supports targeted taxonomic verification and revision. Combining adaptive barcode lengths in one analytical framework offers a general design principle for long-read reference-library construction. Its performance must now be tested in other groups.

## Introduction

Biodiversity science increasingly relies on DNA barcoding to document, monitor and interpret biological diversity across space and time (deWaard et al., 2019; Hebert et al., 2025). Large-scale barcoding and reference library construction increasingly draw on both contemporary field sampling and natural history collections, which together provide voucher-backed material that may span wide variation in molecular integrity (Levesque-Beaudin et al., 2023; Santos et al., 2023). Entomological collections contain billions of specimens, but their use in DNA-based studies requires working with those ranging from recently preserved material to material collected several decades ago (Miller et al., 2013; Shokralla et al., 2011). Fixed-length markers very often create a practical conflict between molecular inclusion and information content. Shorter targets can be recovered more readily from degraded material, whereas longer targets retain more sequence variation and potentially allow greater taxonomic discrimination. Alongside the standard 658 bp cytochrome c oxidase I (COI) Folmer barcode region (Hebert et al., 2010), mini-barcode methods have been established to address this challenge (Meusnier et al., 2008; Prosser et al., 2016; Shokralla et al., 2011).

Earlier COI studies established that shorter targets can extend sequence recovery from degraded specimens (Hajibabaei et al., 2006; Françoso and Arias, 2013; Prosser et al., 2016), while subsequent work evaluated the discriminatory performance of mini-barcodes, classification using shortened queries, and the incorporation of sequences of different lengths in reference databases. However, these components have rarely been evaluated together within a framework that aims to balance preservation-constrained sequence recovery with identification reliability across an order-of-magnitude variation in barcode lengths.

Although generally COI barcoding approaches are successful, COI-based species identification can also be suboptimal (Bensasson, 2001; Leite, 2012; Haran et al., 2015; Cong et al., 2017; Hebert et al., 2023). One reason is the occurrence of non-functional COI copies, termed nuclear mitochondrial DNA segments (NUMTs), which are pseudogenes found in the nuclear genomes of some species and interfere with DNA barcoding (Hedrick, 2011; Hebert et al., 2023). Reported in groups including Orthoptera and Hymenoptera, NUMTs may constitute 0.2% of nuclear genomes (Moulton et al., 2010; Song et al., 2014; Wang et al., 2020; Liu et al., 2024). Although this relationship is not universal (Wang et al., 2020; Ding et al., 2021), NUMT abundance often correlates positively with nuclear genome size (Hebert et al., 2023) and, in Orthoptera, with genomic transposable element content (Liu et al., 2024). Laboratory workflows and bioinformatic filtering can reduce NUMT impacts (Calvignac et al., 2011; Mascolo et al., 2019; Porter and Hajibabaei, 2021; Hebert et al., 2023; Liu et al., 2024), although their negative impact proves inescapable in some groups (Prous et al., 2026). Mitochondrial heteroplasmy can also cause problematic species assignments, although some cases may instead reflect co-amplified NUMTs that are subsequently misinterpreted as heteroplasmy (Parr et al., 2006; Cihlar et al., 2020; Parakatselaki and Ladoukakis, 2021). In sawflies (Symphyta), multiple intra-individual full-length COI-like sequences, many lacking stop codons or length variation, further complicate discrimination between NUMTs and genuine mitochondrial variants (Prous et al., 2026).

Despite such complications, COI barcoding has proven very effective in many hymenopteran lineages (Schmidt et al., 2015; Villalta et al., 2021; Gonçalves et al., 2022), including the most species-rich groups, such as Braconidae and Ichneumonidae (Quicke et al., 2012; Alex Smith et al., 2013). Broad bee (Apiformes) datasets likewise show that COI often performs well overall (Magnacca and Brown, 2012), despite the frequent occurrence of pseudogenes (Creedy et al., 2020). However, several problems have been reported in specific bee lineages, such as co-amplification of non-target DNA, including that of Wolbachia or parasites (Magnacca and Brown, 2012; Bleidorn and Henze, 2021). Primer mismatches and related amplification problems have been reported as potential obstacles in *Andrena*, *Hylaeus* and *Dasypoda* (Magnacca and Brown, 2012; Villalta et al., 2021). Described as a “nightmare taxon” for DNA barcoding, *Dialictus* (*Lasioglossum,* Halictidae) showed a 20% error rate under best-close-match COI identification, with 43 of 110 species occurring in Barcode Index Numbers (BINs) shared with other species (Gibbs, 2018). Heteroplasmy has been reported in *Andrena*, *Bombus* and *Hylaeus* (Magnacca and Brown, 2010, 2012; Ricardo et al., 2020), and stingless bees (Apinae: Meliponini), including *Trigonisca* and *Tetragonula* (Françoso et al., 2023a, 2023b; Yong et al., 2024). A reported lack of COI differentiation among closely related species suggests limited marker resolution in *Sphecodes* (Halictidae), *Scaptotrigona* (Meliponini), and *Colletes* (Colletidae) (Kuhlmann et al., 2007; Magnacca and Brown, 2012; Hurtado-Burillo et al., 2013). NUMTs were also considered problematic in Bombus and Tetragonula (Françoso et al., 2019; Ding et al., 2021). Although combining COI sequences with morphological knowledge helped resolve many problematic *Bombus* cases (Williams et al., 2023, 2024), this approach remains highly labor-intensive and is not readily standardized. Euglossini, a tribe in the subfamily Apinae, shows mixed COI-barcoding performance (Gonçalves et al., 2022). Comparable evidence is scarcer within Euglossa as molecular studies with broad within-species sampling remain uncommon (Ramírez et al., 2010). Nevertheless, COI haplotypes of the morphologically distinct *Euglossa mixta* and *E*. *cognata* have been reported to be intermingled (Dick et al., 2004). The large genome size of up to 3.3 Gb in *Euglossa dilemma* and high transposable element content may promote NUMT prevalence (Brand et al., 2017). NUMT abundance has so far impeded complete mitochondrial-genome assembly (Brand et al., 2017).

The increasingly well-known orchid bees, Euglossini, include approximately 250 described species assigned to five genera: *Aglae*, *Eufriesea*, *Euglossa*, *Eulaema*, and *Exaerete* (Bánki et al., 2025). Orchid bees are key pollinators of not just orchids, and a single species has been recorded to interact with 56 plant families (Pemberton, 2023). Their taxonomy is characterized by ongoing discoveries and revisions, with few molecular studies available (Ramírez et al., 2010; Ayala et al., 2022; Sandoval-Arango et al., 2023). Morphological identification can depend on minute characters that are better documented in males, while females can be difficult to identify or even to associate with conspecific males (Kimsey, 1982; Bembé, 2005). A comprehensive species-level key for the entire tribe does not exist. Several color-based species boundaries remain uncertain because color phenotypes do not always correspond to independently evolving lineages (Ferrari and Melo, 2014; Sandoval-Arango et al., 2023). Even so, broader phylogenetic work indicates that much of the tribe’s taxonomic framework is stable enough for barcode evaluation (Ramírez et al., 2010). Complementing existing DNA barcoding methodologies for Euglossini could significantly enhance tropical research and be applied to other challenging insect genera.

For groups where COI does not deliver species resolution, nuclear ribosomal DNA has been discussed as an alternative (Yao et al., 2010). Its internal transcribed spacer (ITS) region and its components (ITS1, ITS2), have been proposed as a complementary DNA barcode in animals, building on its extensive use in fungi and plants (Yao et al., 2010; Wang et al., 2015). Large-scale comparative analyses across eukaryotes indicate that both ITS1 and ITS2 often provide high success rates in discriminating species (Wang et al., 2015; Yao et al., 2010). As a biparentally inherited non-coding nuclear marker ITS is unaffected by NUMTs or endosymbionts (Klopfstein et al., 2016) and can provide an independent and complementary line of evidence in cases where mitochondrial barcodes are ambiguous. While this comes at the cost of added complexity owing to potential intraspecific and intragenomic variation (Fagan-Jeffries et al., 2019; Marrelli et al., 2006), its utility has been demonstrated in studies on diverse animal lineages including Thysanoptera (Yeh et al., 2015), Malacostraca (Park and Poulin, 2022), Arachnida (Lv et al., 2014; Pérez-Sayas et al., 2022), Collembola (Anslan and Tedersoo, 2015), Copepoda (Bakmaz et al., 2025), Gastropoda (Garzia et al., 2021), Insecta: Diptera (Haarto and Stahls, 2014), Hemiptera (Gomez-Polo et al., 2013), Hymenoptera (Fagan-Jeffries et al., 2019; Ferrer-Suay et al., 2018; Klopfstein et al., 2016). One advantage of the nuclear ribosomal DNA (nrDNA) region is its potential for amplification across a wide range of fragment lengths due to the highly conserved 18S, 5.8S and 28S genes. These provide stable priming targets interspersed with variable regions, such as ITS (Figure 1). The nrDNA system as a species-identifying barcode method therefore permits longer amplicons to be targeted whenever DNA quality allows (Bista and Lino, 2025).

**Figure 1:**
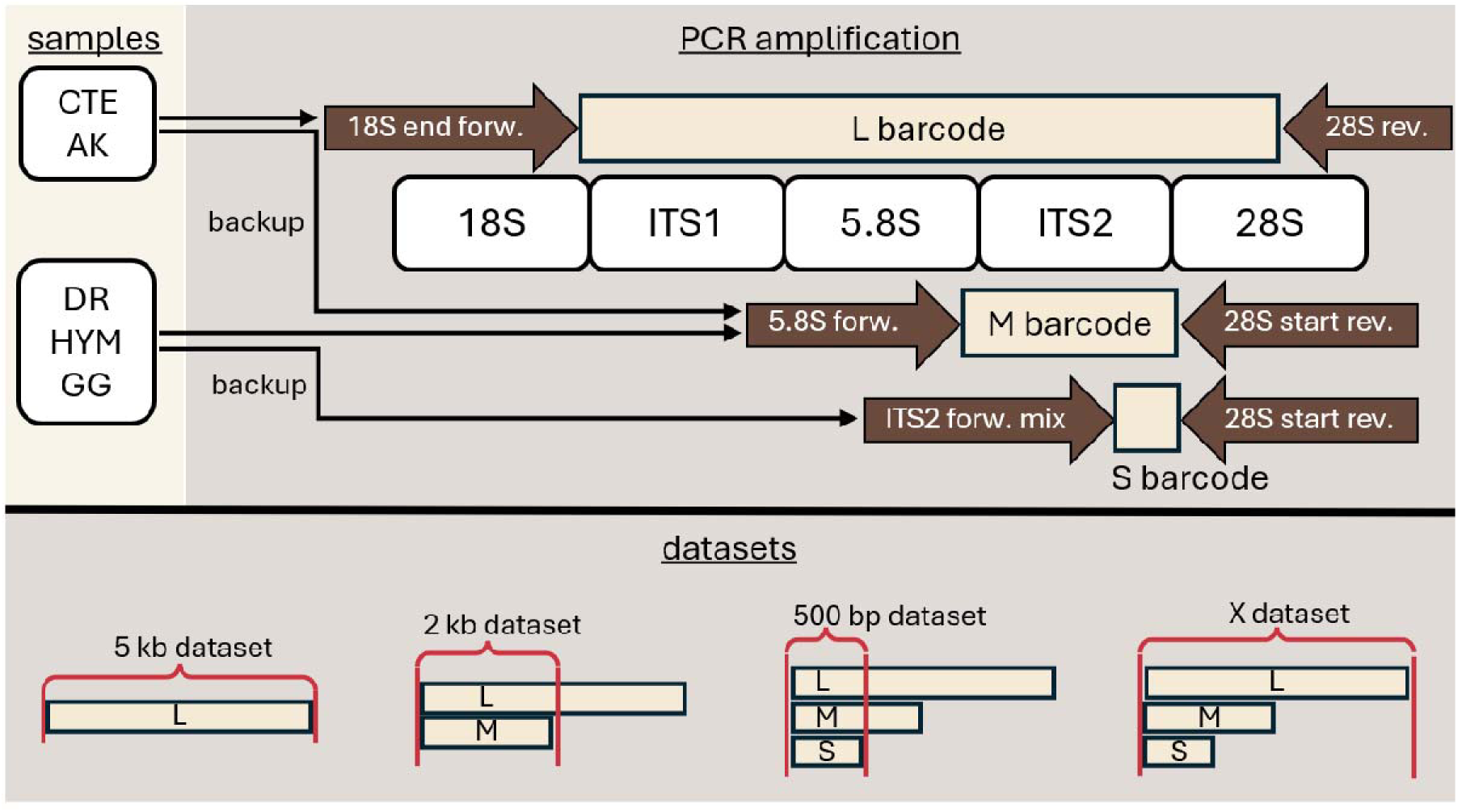
Adaptive-length nrDNA amplification strategy and barcode dataset construction. Sample processing with three different PCR strategies generated overlapping barcode fragments of variable lengths across the nrDNA transcription unit (top panel). The L barcode (yellow box) spans from the end of 18S into the 28S region (white boxes), the M barcode is amplified by the ‘5.8S forward’ and ‘28S start reverse primer’ (brown arrows), and the S barcode (yellow box) overlaps with approximately the last quarter of the M barcode. Barcode datasets (bottom panel), based on L, M and S barcodes (yellow boxes), used for downstream analyses, are named by their approximate length. Longer barcodes are truncated to the same coordinate window to match the respective length class (red bracket), except in the X dataset, which includes all sequences.

In this study, we aimed to develop an adaptive-length nrDNA barcoding framework that links overlapping markers spanning approximately 500 to 5500 bp within a shared analytical workflow and reference library, using Euglossini as a model group. We examined barcode lengths among differing specimens with varied preservation histories and how barcode length affected species discrimination and identification. We further tested whether barcode lengths remained compatible within a shared reference library and how classifier confidence affected the balance between identifying represented species and avoiding erroneous assignments (also termed overclassification) when species were absent from the reference library.

## Methods

### DNA Reference Database Material

To generate the reference library used to evaluate the adaptive-length nrDNA barcoding framework, DNA was extracted from one leg of 590 Euglossini specimens ranging in age from 0 to 31 years (Supplementary Material 1a). These specimens represented all five Euglossini genera: *Aglae*, *Eufriesea*, *Euglossa*, *Eulaema* and *Exaerete*. Two additional Bombus specimens were included as outgroups, giving a total of 592 specimens representing 114 nominal species. Species representation ranged from 1 to 37 specimens per species (mean: 5.1, median: 3). Specimens originated from 13 countries and territories: French Guiana (France) (121), Peru (106), Ecuador (98), Panama (83), Costa Rica (77), Colombia (47), Mexico (33), Guyana (11), Suriname (8), Venezuela (4), Germany (2 *Bombus*), Brazil (1) and Bolivia (1). Specimens derived from five sources: 1) ∼380 from the Collection Thomas Eltz (CTE: TE, CAH and JH series, freshly frozen in alcohol), 2) ∼100 provided by Mabel Alvarado from the Museo de Historia Natural of the Universidad Nacional Mayor de San Marcos in Lima, Peru (HYM, air-dried at ∼45°C), 3) ∼95 from the personal collection of David W. Roubik (DR, air-dried), 4) ∼10 specimens provided by Günter Gerlach (GG, air-dried), and 5) 6 morphologically identified specimens from field collections (AK, freshly frozen). Morphological identifications were provided by multiple experts (Supplementary Material 1a). Targeted morphological re-examinations of problematic specimens were performed on request by multiple specialists, including Thomas Eltz, David W. Roubik, Mabel Alvarado, Thomas Wiesner, Claus Rasmussen and Benjamin Bembé.

Per morphologically identified specimen, one front leg was sampled. For specimens that had previously been partly sampled, the remaining half of the leg was used. Tissue was disrupted in a 1000-µl 96-well Eppendorf plate using one 5-mm stainless-steel ball and two approximately 2-mm AISI 316 steel balls. Samples were processed in a Retsch MM400 mixer mill for two 30-s cycles at 28 Hz. DNA was extracted using a downscaled Mag-Bind® Plant DNA DS 96 Kit (Omega Bio-Tek, cat. no. M1130-01), with all reagent volumes reduced by 50% to minimize costs.

### Molecular Protocols

Primer design began by extracting a complete nrDNA sequence from the chromosome assembly of *Bombus sylvestris* (GenBank OU443152.1). This sequence was used as a BLASTn query to recover nrDNA regions from annotated complete chromosomes spanning Streptophyta, Nemertea, Mollusca, Annelida, Arthropoda and Chordata (Supplementary Material 1b). The resulting set included 33 Apoidea sequences, none from Euglossini. ITS regions were delimited using ITSx, and the flanking 18S, 5.8S and 28S regions were aligned using MAFFT to identify conserved primer binding sites within Hymenoptera (Bengtsson-Palme et al., 2013; Katoh and Standley, 2013). Candidate primers were selected to maximize conservation across hymenopteran sequences while differing from non-hymenopteran taxa, with the selected binding sites conserved across nearly all 33 available Apoidea sequences. However, because no Euglossini sequences were available during primer design, their performance in Euglossini was confirmed empirically. Because no conserved internal priming site could be identified within ITS2, the S barcode primers (Figure 1) were designed as a primer mix, with all primers targeting the same position within ITS2, based on L barcode sequences generated during the first sequencing round. In total, the following primers were designed: 18S forward (18S_end_Ecdysozoa_F), 5.8S forward (58S_end_Hym_F), ITS2 forward (ExEu_ITS2_1522F_PB1F, Euf_ITS2_1522F_PB1F, Eug_ITS2_1522F_PB1F and EugR_ITS2_1522F_PB1F), and 28S reverse (28S_start_Hym_R and 28S_end_Apocrita_R) (Supplementary Material 1c).

A tiered set of three overlapping polymerase chain reaction (PCR) protocols was established, allowing barcode length to be selected according to specimen preservation and expected DNA quality (Supplementary Material 1a). A short barcode (S barcode) recovered a partial ITS2 sequence (∼500 bp) with primers located in the second half of ITS2 and at the 5’ terminus of the 28S sequence (Figure 1). The medium barcode (M barcode) targeted the full ITS2 sequence (∼2000 bp), using the same reverse primer and a forward primer in 5.8S (Figure 1). The long barcode (L barcode) had a median amplicon length of approximately 7100 bp before removal of ITS1 tandem repeats and recovered the full ITS1, ITS2 and the 28S divergent domains D1 to D6 (Gillespie et al., 2005, 2006). After repeat removal, the curated L barcodes used in the analyses had a median length of 5634 bp. DNA extracts from specimens frozen in ethanol (CTE), or from freshly frozen tissue (AK), were amplified using the L barcode protocol with the M barcode protocol as a backup (Figure 1). Air-dried material (HYM, DR and GG) was amplified with the M barcode protocol and employed the S barcode protocol as a backup (Figure 1).

Library preparation used the Oxford Nanopore Technologies Ligation Sequencing Kit SQK-LSK114. Cleanup and size selection were adjusted according to barcode length. L barcode libraries used Long Fragment Buffer (LFB) and a 0.75× bead ratio, while M barcode libraries used Short Fragment Buffer (SFB) and a 0.85× bead ratio. Libraries were sequenced on R10.4.1 flow cells using PromethION P2 Solo (FLO-PRO114M) and MinION (FLO-MIN114) devices. Detailed recommendations for how to adapt and replicate the workflow to other target taxa are available (Supplementary Material 1d).

### Analytical Protocols

Basecalling with dorado v0.8.0 (model: dna_r10.4.1_e8.2_400bps_sup@v4.3.0) was followed by amplicon size filtering (S: 400 bp – 1000 bp, M: 1000 bp – 3000 bp, L: 3000 bp – 15000 bp). A custom R script detected demultiplexing tags and primer sequences with mismatch tolerance, oriented and demultiplexed the reads, and trimmed the primers. Within each sample, reads were pre-clustered at 85% identity, and clusters containing fewer than three reads were removed. All reads from the retained clusters were then returned to the downstream processing pool. Sequence processing included scanning for conserved Euglossini motifs in the L barcode (Supplementary Material 1e). Tandem repeats in the ITS1 region were removed using Tandem Repeat Finder (Benson, 1999). Subsequent analytical steps were performed using the RAMBO pipeline (Kolter and Hebert, 2025). Consensus sequences with more than 10 ambiguous International Union of Pure and Applied Chemistry (IUPAC) nucleotide characters were removed. Where a sample was successfully recovered in two barcode runs (Figure 1), only the longer barcode was retained. Where a specimen produced more than one barcode consensus sequence, the nrDNA variants were preserved and included in the output. Although no clear evidence of pseudogenes remained after initial length filtering, secondary cluster consensus sequences were retained only when their lengths differed by no more than 10% from the dominant cluster consensus. More divergent sequences were excluded as a precaution because genuine nrDNA variants within a specimen are not expected to differ by more than 10% after tandem-repeat removal.

To separate marker performance from taxonomic uncertainty, sequence records were screened for taxonomic inconsistencies before classifier validation. Potential conflicts were flagged using a maximum-likelihood phylogeny inferred from the nrDNA alignment with IQ-TREE 3 (Wong et al., 2026) under automatic model selection and 1000 bootstrap replicates (Supplementary Material 1f). Sequences falling outside their expected subgenus, sensu Ramírez *et al*. (2010), were re-evaluated using independent evidence, including a re-assessment of existing morphological identifications and discussions in the taxonomic literature. Where supported, revised taxonomic assignments were incorporated before validation (Supplementary Material 1a). In a small number of cases, closely related species were treated jointly as species groups (see Discussion).

Three barcode-length datasets were constructed from the curated consensus sequences for the length-specific analyses. The 5 kb dataset comprised only L barcodes (Figure 1). The 2 kb dataset contained all M barcodes and L barcodes truncated to the same conserved flanking region, thereby covering the entire ITS2 region. The 500 bp dataset comprised the S barcodes plus the M and L barcodes trimmed to the same region (Figure 1).

To evaluate species separation, an indicator of DNA barcoding performance, pairwise sequence dissimilarities were calculated within each barcode-length dataset using two metrics (Figure 2). First, alignment-based mismatches were calculated using an IUPAC-aware Hamming metric (Ullrich, 2025). Second, an inverse-document-frequency (IDF)-weighted k-mer dissimilarity was calculated using IDTAXA in the DECIPHER R package (Wright, 2025). This weighting reduces the contribution of common k-mers shared across many sequences and is hereafter referred to as the IDTAXA k-mer distance. Intragenomic comparisons involved RAMBO cluster consensus variants from the same specimen, intraspecific comparisons involved different specimens of the same species, and congeneric interspecific comparisons involved different species within the same genus. Two forms of separation were evaluated for each sequence. The nearest-neighbor analysis compared, for each sequence, the minimum distance to a conspecific sequence from another specimen with the minimum distance to a congeneric heterospecific sequence. The DNA barcoding-gap analysis compared the maximum intraspecific distance with the minimum congeneric interspecific distance (Meyer and Paulay, 2005). For both comparisons, separation was expressed as the normalized distance difference (*d*_inter_ − *d*_intra_) / (*d*_inter_ + *d*_intra_), where *d_intra_* represents the relevant minimum or maximum intraspecific distance. Positive values indicate positive separation, while values of zero or below indicate ties or reversals (Figure 2).

**Figure 2:**
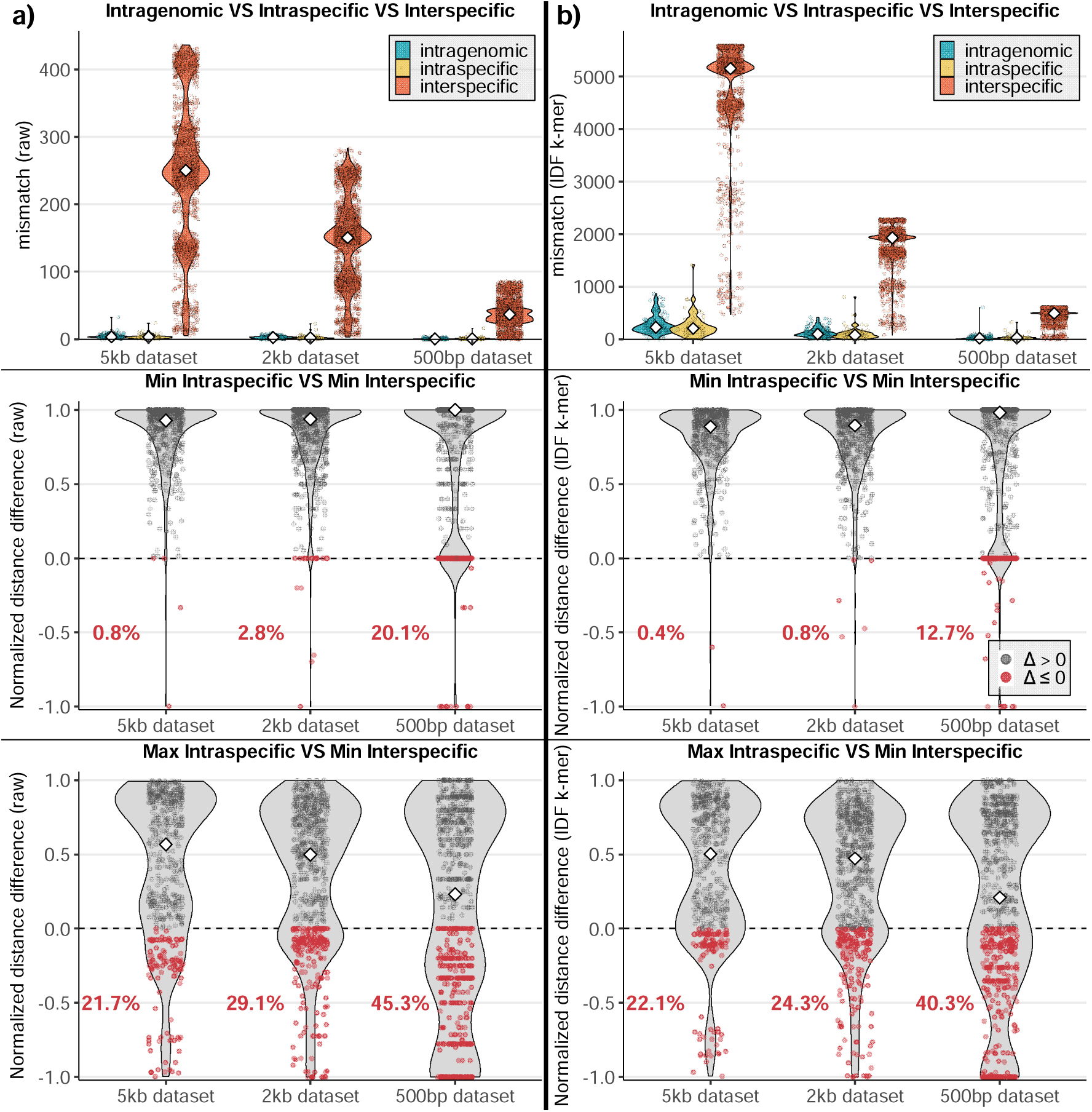
Pairwise mismatch structure across datasets and distance metrics. Columns show the IUPAC-aware Hamming distance in (a) and an IDF-weighted k-mer Jaccard dissimilarity which was calculated from k-mers and inverse-document-frequency weights for (b) for the 5 kb, 2 kb, and 500 bp datasets. The top row shows pairwise intragenomic (blue), intraspecific (yellow), and interspecific (orange) distances within genera. In the top row, intragenomic points represent specimen medians, intraspecific points represent species medians, and interspecific points represent randomly sampled congeneric interspecific sequence pairs. Distances are shown as raw mismatches in (a) and IDF-weighted k-mer distances in (b). These scales are not directly comparable. The middle row shows, for each sequence, the normalized distance difference between its minimum intraspecific and minimum interspecific distances. The bottom row shows the normalized distance difference between the maximum intraspecific and minimum interspecific distances. Normalized distance differences were calculated as (*d*_inter_ − *d*_intra_) / (*d*_inter_ + *d*_intra_). In the middle and bottom rows, each point represents one sequence. Positive values (grey) indicate positive separation, whereas values of zero or below (red) indicate ties or reversals. Percentages indicate the proportion of sequences with values of zero or below.

Subsequent classification analyses were conducted with IDTAXA within a leave-k-out validation framework over 16 rounds. In each round, at most one query specimen was selected from each species represented by at least two specimens. All consensus sequences derived from the selected query specimen were removed from the training set. All remaining sequences were retained in the reference set for classifier training. Across rounds, each specimen was used as a query at most once. Thus, the 16 rounds fully covered all eligible specimens for 62 of 66 analysis taxa in the L dataset, 82 of 87 in the M dataset, and 86 of 91 in the S dataset. Taxa represented by more than 16 eligible specimens were subsampled, resulting in 320, 455, and 519 tested specimens, respectively. The four species with subsampled coverage were *E. Orellana* (37 specimens), *E. ignita* (31 specimens), *E. decorata* (24 specimens), and *E. hemichlora* (23 specimens). Because each query species remained represented in the reference set during every round, this setup is here referred to as the reference-present validation.

For the reference-present validation, all classification outcomes were summarized at the specimen level. Assignment rate was defined as the proportion of query specimens receiving a species-level assignment. Correct-identification rate was the proportion of query specimens assigned to the correct species. Precision was the proportion of assigned specimens that were assigned to the correct species.

A second, more stringent benchmark was implemented as a reference-absent validation. At each IDTAXA threshold, 100 independent omission rounds were performed. In each round, 20% of species represented by at least three specimens were randomly removed from the reference library, and all specimens belonging to these taxa were used as queries. Because the query taxa were absent from the classifier database, correct species-level assignments were impossible by design. Overclassification was therefore defined as the proportion of query specimens receiving a species-level assignment despite the absence of their taxon from the reference library and was analyzed separately from reference-present correct-identification rates (Figure 3).

**Figure 3:**
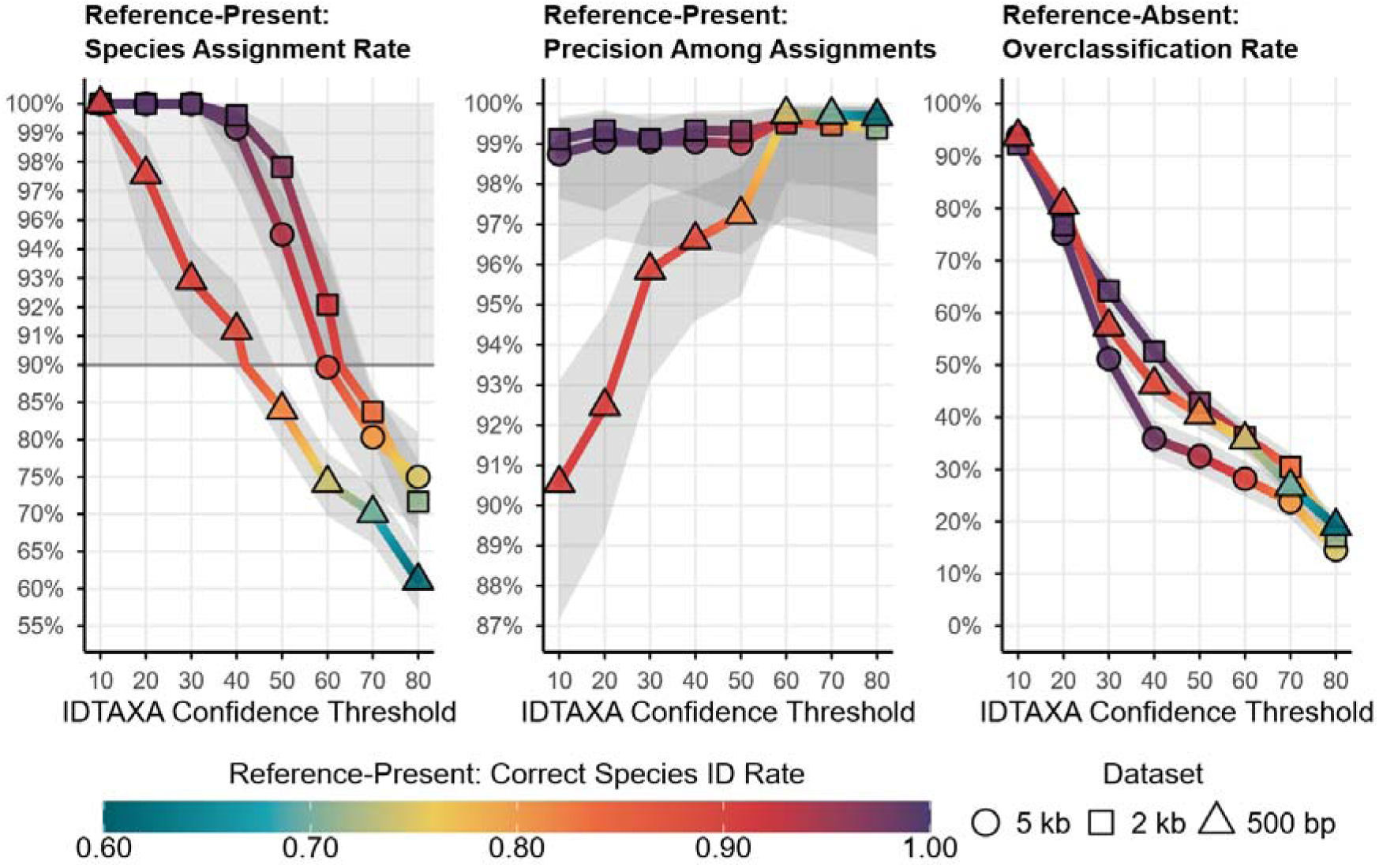
Performance of the IDTAXA classifier across confidence thresholds and sequence datasets Performance of the IDTAXA taxonomic classifier evaluated at the specimen level across increasing confidence thresholds for three marker datasets (shape). Panels show the reference-present assignment rate (left), reference-present precision among accepted assignments (center), and reference-absent overclassification rate (right) as functions of the IDTAXA confidence threshold. Grey bands indicate 95% confidence intervals around the pooled reference-present rates, clustered across 16 leave-k-out rounds, and the mean reference-absent rates across 100 omission rounds. Reference-present correct species identification rate (color) was calculated as the product of assignment rate and precision among accepted assignments, did not consider overclassification, and was applied consistently across all panels. Values below 0.75 were present in the data but set to the minimum of the color scale. For the assignment rate panel, the upper range between 0.9 and 1.0 is visually expanded to enhance discrimination among high-performing thresholds (shaded background). The precision panel is displayed on a linear scale restricted to the high-performance range. Values are derived from leave-k-out cross-validation across all datasets (except the mixed-length X dataset) and thresholds. Because the three marker-length datasets differ in taxonomic and specimen composition, the figure compares the performance of the complete marker-specific reference datasets.

IDTAXA threshold values were assessed within both validation schemes. In IDTAXA, the threshold specifies the minimum bootstrap support required to accept an assignment at a given taxonomic rank. Otherwise, the query is reported as unclassified at that rank. IDTAXA confidence thresholds from 10 to 80 were tested in increments of 10, with 250 bootstrap replicates. Dataset-specific working thresholds were selected at the breakpoint where further reductions in reference-absent overclassification were accompanied by disproportionately larger losses in reference-present correct-identification rate.

Barcode interoperability across marker lengths was then evaluated using the same reference-present leave-k-out framework in two additional analyses. First, leave-k-out validation was performed on the mixed-length X dataset (Figure 1), in which both query and reference sets contained L, M, and S barcodes. Second, interoperability was assessed by using the 2 kb dataset, which comprised M barcodes and truncated L barcodes, as the query set, and the 5 kb dataset, containing only full-length L barcodes, as the training reference for IDTAXA. In this second analysis, when a selected 2 kb query corresponded to a truncated L barcode, all full-length L sequences from the same specimen were removed from the 5 kb training set to ensure that the query specimen was not represented in both sets.

## Results

Recoverable barcode length varied with specimen preservation history. Amplification success of the L barcode exceeded 95% for specimens preserved in 95% ethanol within hours of collection and stored at −20°C. Specimens yielding L barcodes were 9.2 ± 6.2 years old (mean ± standard deviation, n = 366). Under this preservation regime, amplicons of up to 8000 bp were obtained. TE8, collected in 2008, was the oldest specimen provided for this study under this preservation regime and yielded an L barcode after more than 15 years of storage. Its successful amplification indicates that the maximum storage duration permitting L barcode recovery was not reached within the examined material.

For air-dried material, drying temperatures generally did not exceed 45°C, although the exact conditions were unknown in some cases. The M barcode was attempted first, with the S barcode used when M barcode amplification was unsuccessful. Specimens yielding M barcodes were 3.5 ± 4.2 years old (n = 154). Of the 95 HYM specimens, 34 yielded M barcodes, whereas 61 yielded only S barcodes. Specimens yielding S barcodes were on average 11.7 ± 6.5 years old. The oldest successful S barcode specimen was the 31-year-old GG0602, collected in 1993.

Before tandem-repeat removal, L barcode amplicons showed substantial length variation, with a median length of 7137 bp (interquartile range (IQR): 569 bp). This variation corresponded to differences in tandem-repeat copy number. Repeat units had a mean length of 221 bp, and amplicons contained a median of four repeats (IQR: 2–7). After removal of the repeat region, L barcode consensus sequences had a median length of 5634 bp and an IQR of 79 bp (∼1.4%). M barcodes had a median length of 2064 bp with an IQR of 56 bp (∼2.7%), whereas S barcodes had a median length of 513 bp with an IQR of 38 bp (∼7.4%). Ambiguous IUPAC nucleotides were rare, accounting for 0.036% of nucleotides in L barcodes, 0.084% in M barcodes, and 0.116% in S barcodes. Overall, a mean of 1.4 cluster consensus sequences was retained per specimen, potentially representing intragenomic barcode variants (Supplementary Material 1g).

The main analytical pipeline focused on the 5 kb, 2 kb, and 500 bp datasets, which contained 366 specimens representing 83 taxa, 520 specimens representing 108 taxa, and 590 specimens representing 114 taxa (Figure 1), respectively (Supplementary Material 1h).

We examined pairwise genetic distance distributions within and between taxa under the curated taxonomic framework described in Methods. These distance distributions were assessed using two metrics: an IUPAC-aware Hamming mismatch distance (Figure 2a) and an IDF-weighted k-mer score (Figure 2b). Across all pairwise sequence-distance summaries (Figure 2, top), both metrics showed the same overall trend, with interspecific values generally exceeding intragenomic and intraspecific values. The median interspecific value exceeded the mean of the intragenomic and intraspecific medians by ∼70-fold under the Hamming metric across all marker datasets and by ∼24- to 38-fold under the IDF-weighted k-mer metric. Dispersion of the interspecific distributions differed strongly between metrics, with interspecific IQRs corresponding to 48% to 60% of the median under the Hamming metric (5 kb: 48%, 2 kb: 60%, 500 bp: 52.6%), compared with 15.2% to 18.5% under the IDF-weighted k-mer metric (5 kb: 15.2%, 2 kb: 15.8%, 500 bp: 18.5%). Across all three datasets, intragenomic and intraspecific mismatch distributions were highly similar, with consistently high overlap coefficients (0.932–0.941) and low Hellinger distances (0.112–0.164), indicating that the central mass of both distributions was largely shared.

In the nearest-neighbor analysis (Figure 2, middle), ties or reversals were recorded when the nearest congeneric heterospecific sequence was as close as or closer than the nearest conspecific sequence from another specimen. These cases were uncommon in the 5 kb and 2 kb datasets under both metrics but increased sharply in the 500 bp dataset, affecting 20.6% of sequences under the IUPAC-aware Hamming metric and 12.9% under the IDF-weighted k-mer metric.

In the DNA barcoding gap analysis (Figure 2, bottom), the proportion of sequences lacking positive separation between the maximum intraspecific and minimum congeneric interspecific distances increased as marker length decreased under both metrics. In the 500 bp dataset, this proportion was 45.3% under the IUPAC-aware Hamming metric and 40.3% under the IDF-weighted k-mer metric. The lower quartile of the normalized distance difference also shifted from −0.20 under the Hamming metric to −0.03 under the IDF-weighted k-mer metric, indicating that failures were both less frequent and less severe under the latter metric.

Given the clearer separation under the IDF-weighted k-mer score, classification performance was subsequently evaluated with IDTAXA. Evaluation across confidence thresholds identified working thresholds of 40 for the 5 kb dataset, 50 for the 2 kb dataset, and 40 for the 500 bp dataset. At these thresholds, correct-identification rates were 98.1%, 96.9%, and 88.2%, respectively, while reference-absent overclassification rates were 35.9%, 42.8%, and 46.3%. Assignment rates remained high across datasets (91.3–99.1%), as did precision (96.6–99.3%), although the 500 bp dataset showed the only notable reduction in precision. Across all datasets, increasing the confidence threshold reduced overclassification at the cost of lower correct-identification rates (Figure 3).

When specimen-level outcomes were first averaged within species and then across species, giving each evaluated species equal weight, correct-identification rates were 98.9%, 98.2%, and 92.2% for the 5 kb, 2 kb, and 500 bp datasets, respectively, compared with pooled specimen-level rates of 98.1%, 96.9%, and 88.2%. The difference was greatest for the 500 bp dataset, indicating that identification failures were concentrated in several comparatively well-represented species. A matched-specimen analysis confirmed that this decline was not caused by differences in query composition. The 81 tested specimens belonging to 18 taxa absent from the 5 kb dataset were correctly identified, whereas all 61 failures in the 500 bp dataset occurred in taxa already represented by L barcodes.

All tested *Euglossa dilemma* specimens in the 5 kb, 2 kb, and 500 bp datasets, respectively, were correctly assigned to the combined *E. viridissima–E. dilemma* analysis group. Scoring these specimens as unsuccessful under their original species designation would reduce pooled specimen-level success to 95.6%, 95.6%, and 86.9%, and species-averaged success to 97.4%, 97.1%, and 91.2%, respectively. Complete identification of all tested specimens was achieved for 62 of 66 taxa in the 5 kb dataset, 80 of 87 taxa in the 2 kb dataset, and 78 of 91 taxa in the 500 bp dataset. No taxon failed completely with the L or M barcode, whereas four taxa failed completely with the S barcode. *Euglossa orellana* showed a particularly clear length-dependent decline, with 14 of 16 specimens correctly identified using the L barcode, 8 of 16 using the M barcode, and none of 16 using the S barcode. Genus-level identification was 100% in all three reference-present validations.

The same leave-k-out cross-validation via IDTAXA applied to the X dataset (Figure 1), comprised of L, M, and S barcodes, achieved a mean assignment rate of ∼98% and a mean precision of ∼98%, yielding a reference-present species identification success of ∼97% at a confidence threshold of 40. These results were based on 462 L, 185 M, and 70 S barcodes in the test sets. The high pooled performance was not attributable to the numerical dominance of longer barcodes, because specimens represented by S barcodes in the X dataset were not a poorly performing minority. When all specimen sequences were cut to the S barcode region, correct-identification rates were 98.4% (62/63), 94.4% (135/143), and 83.4% (261/313) for specimens represented in the X dataset by S, M, and L barcodes, respectively. The L barcode subset included most of the taxonomically difficult groups and accounted for 52 of the 61 failed identifications (85.2%) in the example above. Overclassification was not evaluated for this mixed-length analysis because the reference-absent setup is not directly comparable across barcode lengths.

We also evaluated how sequences from the 2 kb dataset perform when queried against the 5 kb reference dataset to assess the impact of reference length on classification success. For the leave-k-out cross-validation using sequences from the 2 kb dataset as queries versus the 5 kb dataset as training data we chose a more conservative threshold value of 60. 320 samples were tested and the reference-present success of ∼90% was very similar to results of the leave-k-out cross-validation of the 2 kb dataset versus itself at the same threshold, which was ∼92% (Figure 3).

## Discussion

Two current limitations in DNA barcoding are addressed in this study. First, as long-read biodiversity sequencing evolves from proof-of-concept studies to broader applications (Bista and Lino, 2026), the standard 658-bp COI barcode does not systematically exploit the additional sequence information accessible through longer reads. Second, COI provides insufficient species-level resolution in some Hymenoptera, and its use can be complicated by other biological and methodological factors documented across different lineages (Klopfstein et al., 2016; Gibbs, 2018; Prous et al., 2020; Bleidorn and Henze, 2021; Gonçalves et al., 2022; Vasilita et al., 2022; Jafari et al., 2023; Šet et al., 2024). The adaptive-length nrDNA framework presented here addresses both of these limitations by offering an alternative to the standard COI barcode by a length-scalable nrDNA marker that exploits longer reads when available, while simultaneously providing a length-scalable nuclear alternative when problems specific to mitochondrial DNA are a concern. In practical terms, the results show that long-read barcoding and museum-linked sampling do not have to be treated as separate systems. Instead, they can be connected by overlapping markers that preserve analytical compatibility while accommodating differences in molecular integrity.

### Specimen Preservation and Recoverable Barcode Length

As DNA-based identification and sequencing increasingly underpin biodiversity research (Schmid et al., 2025), the future molecular value of natural history collections will be enhanced as systematic tissue banking and preservation methods optimized for DNA integrity become integral components of modern collection practice (Bakker et al., 2020; Eymann et al., 2010; Gostel et al., 2016; Ogiso-Tanaka et al., 2025; Schindel and Cook, 2018; Ståhls et al., 2021). However, established museum preservation practices are developed primarily for morphological study and do not always support long-term DNA-based research (Bakker et al., 2020; Van Caenegem and Haelewaters, 2024), raising concerns about future molecular accessibility as taxonomic expertise declines and molecular data generation accelerates (Coleman and Radulovici, 2020; Kodama et al., 2012; Löbl et al., 2023; Paknia et al., 2015). In the material analyzed here, long amplicons were successfully sequenced from specimens preserved in 95% ethanol stored at −20°C, a pattern that aligns with broader methodological work showing that high-concentration ethanol combined with cold storage best preserves DNA for downstream sequencing (Marquina et al., 2021). Existing collections, however, contain specimens with widely differing preservation histories. Adjusting barcode length to specimen preservation has a long precedent in COI barcoding, where varied fragment lengths have been used to recover sequences from degraded material (Hajibabaei et al., 2006; Meusnier et al., 2008; Shokralla et al., 2011; Françoso and Arias, 2013; Prosser et al., 2016). Similarly, the recovery of the S barcode from older air-dried material demonstrates how a shorter target can extend molecular access. However, by jointly assessing genetic separation, identification success, and overclassification, the present framework extends length adaptation from a means of recovering sequences to a calibrated strategy for managing identification reliability throughout the barcoding workflow. Combined with long-read sequencing, this strategy operates in both directions, uniting shorter fallback barcodes for degraded material with barcodes extending far beyond conventional lengths when DNA integrity permits.

### Sequence Data Analysis and Length-Dependent Differences in Barcode Performance

The recovery of an average of 1.4 consensus clusters per specimen is consistent with the suspected presence of multiple candidate intragenomic nrDNA variants. The extent and consequences of ITS variation differ among Hymenoptera, ranging from little or no detected intraspecific variation to substantial intragenomic variability (Alvarez and Hoy, 2002; Veen et al., 2003; Fagan-Jeffries et al., 2019). The retained variants showed no pseudogene-like sequence characteristics or anomalous phylogenetic placement and they consistently clustered with other sequences of the respective species. They are therefore interpreted as intragenomic nrDNA variants, potentially reflecting incomplete homogenization among repeated nrDNA units (Álvarez and Wendel, 2003; Feliner and Rosselló, 2007). In the present Euglossini dataset, the retention of multiple nrDNA variants was compatible with high reference-present identification success (Figure 3). Performance was evaluated at the specimen rather than sequence level, thereby preventing specimens represented by multiple variants from receiving greater weight.

Barcode-based identification depends not only on the sequence variation provided by a marker, but also on the analytical framework used to interpret that variation (Tanabe and Toju, 2013; Orsholm et al., 2026). This choice becomes particularly important when a species lacks a clear barcoding gap, defined as a minimum interspecific distance exceeding its maximum intraspecific distance (Meyer and Paulay, 2005). Such separation remained incomplete in our nrDNA data (Figure 2), consistent with its limited occurrence in available COI data for Euglossini and other Apidae datasets (Gonçalves et al., 2022; Šet et al., 2024). The extent of the resulting ambiguity, however, depended on the analytical framework, where Hamming and IDF-weighted k-mer comparisons produced similar results for the 5 kb dataset, whereas the k-mer metric reduced ambiguous distance relationships for the 2 kb and particularly the 500 bp dataset (Figure 2). The heterogeneous structure of the nrDNA combines conserved ribosomal RNA genes with rapidly evolving ITS regions, where indel accumulation and residual intragenomic variation complicate alignment and produce an uneven distribution of taxonomically informative variation (Álvarez and Wendel, 2003; Feliner and Rosselló, 2007; Mishra et al., 2021). Hamming distance treats all aligned nucleotide mismatches equally, whereas IDF weighting in the IDTAXA framework downweights k-mers shared across many taxa and emphasizes those with greater discriminatory value (Murali et al., 2018; Wright, 2025). The combination of alignment-free k-mer representation and differential feature weighting may therefore retain more useful taxonomic signal as barcode length decreases, making the framework well suited to an adaptive-length reference database. A recent benchmark provides an independent parallel, with composition-based classifiers, including IDTAXA, performing best for fungal ITS, a marker that is difficult to align reliably but rich in short-motif signal (Orsholm et al., 2026). Although fungal ITS and Euglossini nrDNA are not directly comparable, both contain indel-rich spacer regions with unevenly distributed informative sequence features.

Although distance analyses describe the separation between intra- and interspecific variation, they do not directly establish the reliability of specimen identification (Collins and Cruickshank, 2013). Reference-present validation, in which each test species remained represented in the training data, yielded 88.2–98.1% correct species identification across barcode lengths (Figure 3). This conventional design tests discrimination among represented species, but not whether a classifier recognizes that the query species is absent, a distinct problem termed novel-taxon classification, out-of-distribution detection or simply overclassification (Murali et al., 2018; Fujisawa and Imai, 2026; Orsholm et al., 2026). Without an explicit rejection criterion, nearest-neighbour or highest-identity matching necessarily assigns every reference-absent query to a represented species, resulting in a 100% overclassification. IDTAXA instead uses bootstrap-derived confidence at each taxonomic rank to withhold unsupported assignments (Murali et al., 2018). At the selected thresholds, overclassification was 35.9% in the 5 kb dataset, 42.8% in the 2 kb dataset, and 46.3% in the 500 bp dataset, showing that IDTAXA withheld species-level assignments for most reference-absent specimens but did not eliminate the risk arising from incomplete reference coverage (Figure 3). More importantly, shortening affected the two validation conditions unequally. The 2 kb dataset approached the 5 kb dataset in reference-present identification but were less effective at rejecting unsupported assignments when the species was absent. This divergence agrees with evidence that unknown-species detection is more susceptible than known-species identification to the loss of diagnosable characters in short barcodes (Fujisawa and Imai, 2026). Reference-absent validation therefore reveals that the principal practical advantage of the L barcode over the M barcode lies in its more reliable behaviour under incomplete reference coverage, a condition increasingly recognized as requiring separate evaluation in classifier benchmarks (Murali et al., 2018; Orsholm et al., 2026). Although the 5 kb, 2 kb and 500 bp sequence databases differed in composition, the matched-specimen and taxon-composition controls showed that the lower 500 bp performance was not driven by specimens or taxa added only at shorter barcode lengths. Instead, the failures were concentrated in difficult taxa already represented by L barcodes. This largely reflected the presence of multiple taxonomically difficult groups in the L subset for which barcode discrimination declined after truncation.

The remaining overclassification in the 5 kb dataset suggests limited genetic separation among some closely related species. This interpretation is consistent with literature describing several taxonomically problematic groups in Euglossini as difficult to resolve using both morphological and molecular data (Dressler, 1982; Faria and Melo, 2007; Ramírez et al., 2010; Hinojosa-Díaz and Engel, 2011; Faria and Melo, 2012). These results suggest that DNA barcoding may be of limited value as an independent basis for species delineation in closely related groups with undescribed species, supporting the view that species discovery and specimen identification are separate objectives (Collins and Cruickshank, 2013; Miralles et al., 2024). By contrast, once correctly identified voucher specimens are represented in the reference database, barcoding-based specimen identification can still be highly effective even in these difficult complexes. Within this broader framework, the performance of the L barcode is biologically plausible as longer ribosomal targets provide greater discriminatory power because they sample more informative variation across the cistron (Heeger et al., 2018; Krehenwinkel et al., 2019), while shorter targets remain valuable mainly because they preserve molecular access when DNA quality or specimen history limit recovery of longer amplicons (Little, 2014; Chen et al., 2023). Taken together, our results do not support a single universally optimal barcode, but instead a hierarchical strategy in which L maximizes resolution, M provides a practical compromise, and S extends inclusion of material that would otherwise be lost from a molecular analysis.

### Taxonomic Interpretation and Limits of nrDNA Barcoding

The following cases illustrate how nrDNA data can flag taxonomic inconsistencies and guide targeted re-examination, while interpretation of species boundaries remains dependent on independent morphological, chemical, population-genetic, or published taxonomic evidence.

The reference dataset analyzed here covers 114 of the 252 currently documented species of Euglossini (Bánki et al., 2026) and includes multiple sets of closely related taxa, including nine members of the *Euglossa cordata* species group. To our knowledge, it represents the largest curated collection of Euglossini barcodes derived from morphologically identified specimens. Nevertheless, some taxonomic uncertainty remains, most notably within material provisionally assigned to *E*. *decorata*, which probably includes several described species that could not be identified reliably using the available taxonomic keys (Hinojosa-Díaz and Engel, 2011; Ferrari and Melo, 2014). Although the broader phylogenetic implications of these data will be addressed in follow-up studies, the observed patterns provide several distinct examples of how nrDNA results can be interpreted in relation to existing taxonomic evidence.

1. Specimens identified as *Euglossa crassipunctata* were split into two clusters, labeled here as *E. crassipunctata* 1 and 2. Literature describes *E*. *clausi* and *E*. *moratoi* as close relatives of *E*. *crassipunctata* and documented that specimens of at least *E*. *clausi* had previously been treated as *E*. *crassipunctata* (Nemésio and Engel, 2012). This provides taxonomic precedent for heterogeneity among specimens assigned this name, but does not determine the identity or taxonomic status of either nrDNA cluster.
2. *Euglossa ignita* was split into three clusters labelled as *E*. *ignita* 1, 2 and 3. Previous molecular work had already recovered divergent and geographically unstructured COI haplotypes within *E*. *ignita*. Three intraspecific clades formed an unresolved polytomy with *E*. *flammea*, *E*. *orellana*, and *E*. *chalybeata*, which was interpreted as evidence of recent speciation and retention of ancestral alleles (Dick et al., 2004). At least two of the evident *E*. *ignita* lineages were found to be separable by morphological and chemical (perfumes) characters (Eltz et al., in preparation). Taken together, these patterns suggest that nrDNA barcoding can help reveal recent divergences and previously missed lineage structure within taxonomically difficult Euglossini groups.
3. Other cases revealed conflicts rather than corroborated lineage structure. In our dataset, *E. iopyrrha* is nested within *E. mixta*. This contrasts with earlier reports placing *E. iopyrrha* closer to *E. cognata*, a species that is otherwise molecularly distinct from *E. mixta* (Ramírez et al., 2010). We confirmed repeatedly by morphology that our specimens identified as *E*. *iopyrrha* clearly showed the diagnostic red-bronze fourth tergite (Dressler, 1978). Recurring identification problems were described for Atlantic Forest specimens previously identified as E. *iopyrrha* and *E*. *mixta*, which ultimately represented the distinct species *E*. *botocuda* and E. *calycina*, respectively (Faria and Melo, 2012). The observed nesting revealed by the nrDNA barcodes warrants expanded specimen collection and sequencing to clarify the taxonomic status of the investigated specimens.
4. For classifier evaluation, *E*. *obtusa* and *E*. *dodsoni* were treated as a single taxon because the nrDNA barcodes did not support their reliable separation. Earlier suggestions indicate that *E*. *obtusa* “ may eventually prove to be a subspecies of *E*. *dodsoni*“ (Dressler, 1978). In addition, a high degree of similarity between both species has been described previously in the literature (Hinojosa-Díaz and Engel, 2011). The nrDNA barcode data suggest a taxonomic validation of these two species might be a revealing new insight.
5. *Euglossa dilemma* and *E. viridissima* represent an exceptional case within *Euglossa* (Eltz et al., 2011). The two are clearly distinct species based on population-genetic evidence from single-nucleotide polymorphisms, differences in morphology and pheromone analogues (also known as male ‘perfumes’), and evidence of reproductive isolation across a broad zone of sympatry (Brand et al., 2020). However, their relatively recent divergence time of approximately 150 000 years seems not to have produced sufficient DNA sequence differences for reliable species-level separation by an analysis based either on COI barcodes or the nrDNA barcode data (Eltz et al., 2011). They were treated as a single taxon for classifier evaluation to enable assignment at the highest resolution supported by the nrDNA marker, rather than forcing a fallback to genus level. This analytical treatment does not, however, challenge their recognition as separate species.

Taken together, these case studies illustrate three distinct outcomes of nrDNA barcoding: corroboration of independently supported lineage structure, detection of conflicts requiring taxonomic re-examination, and failure to distinguish the youngest documented species of *Euglossa*, widely supported by other evidence. Such molecular flagging is particularly valuable in Euglossini, where species identification faces challenges, and speciation by rapid sorting into ‘perfume groups’ essential to reproductive integrity may hasten population divergence (Bembé, 2005; Faria and Melo, 2007; Ramírez et al., 2010; Hinojosa-Díaz and Engel, 2011). Beyond improving specimen identification, the present dataset is useful by prioritizing particular taxa and specimens for targeted follow-up. This role is consistent with the broader use of barcode and genomic evidence in insect systematics to expose cryptic diversity and organize taxonomic re-evaluation, while emphasizing that molecular evidence is most informative when interpreted together with independent evidence rather than as stand-alone proof of species boundaries (Schlick-Steiner et al., 2010; Padial et al., 2010; Miller et al., 2016).

## Conclusion

The central implication of this study is that DNA barcoding need not be organized around a single fixed marker length. By linking overlapping nrDNA targets within one analytical framework, specimens of differing molecular integrity can contribute to the same reference system while retaining as much sequence information as their preservation state allows. In this way, degradation changes the amount of recoverable information rather than determining whether a specimen can participate in the analysis at all. For taxonomically difficult groups such as Euglossini, this provides a practical route to integrate recently collected and historical material while explicitly accounting for limitations arising from marker length, incomplete reference databases, and uncertain species separation. More broadly, the adaptive-length design offers a general principle for long-read barcoding: recover the longest informative target supported by each specimen and also preserve analytical interoperability across marker lengths.

## Funding Statement and Competing Interests

Funding was provided by the Gordon and Betty Moore Foundation (Andes Amazon Fund). The authors declare no competing interests, financially or otherwise.

## Data Accessibility Statement and Acknowledgements

Sequences were deposited in the European Nucleotide Archive (ENA) study PRJEB106079, while supplementary raw data are available at Zenodo (doi:10.5281/zenodo.18237409).

We gratefully acknowledge Thomas Wiesner and Benjamin Bembé for supporting the morphological specimen verification step. Insects from the Museo de Historia Natural (Universidad Nacional Mayor de San Marcos, Lima, Peru) were exported with permission N°003858-SERFOR, issued by the Servicio Nacional Forestal y de Fauna Silvestre.

An AI-based language and coding assistance tool (ChatGPT 5.6) was used during manuscript preparation to support language editing and code review. All analytical decisions, code implementation, validation, and interpretation of results were performed and verified by the authors.

**Andreas Kolter:** Conceptualization, Methodology, Software, Validation, Formal analysis, Investigation, Data curation, Writing – original draft, Writing – review and editing, Project administration, Visualization.

**Thomas Eltz:** Resources, Investigation, Validation, Writing – review and editing.

**Mabel Alvarado:** Resources, Investigation, Validation, Writing – review and editing.

**David W. Roubik:** Resources, Investigation, Validation, Writing – review and editing

## Notes

### Competing Interest Statement

The authors have declared no competing interest.

